# Genomics-based deconvolution of multiplexed transcriptional responses to wood smoke particles defines rapid AHR signaling dynamics

**DOI:** 10.1101/2021.02.23.432311

**Authors:** Arnav Gupta, Sarah K Sasse, Margaret A. Gruca, Lynn Sanford, Robin D. Dowell, Anthony N. Gerber

**Author notes:** To whom correspondence should be addressed: Dr. Anthony N. Gerber, Department of Medicine, National Jewish Health, Room K729, 1400 Jackson St, Denver, CO, 80206, USA.

## Abstract

Heterogeneity of respirable particulates and compounds complicates understanding transcriptional responses to air pollution. Here, we address this problem through applying precision nuclear run-on sequencing (PRO-seq) to measure nascent transcription and the assay for transposase-accessible chromatin using sequencing (ATAC-seq) to airway epithelial cells after wood smoke particle (WSP) exposure. We used transcription factor enrichment analysis to agnostically identify temporally distinct roles for the TCF/SRF family, the aryl hydrocarbon receptor (AHR), and NFkB in regulating transcriptional changes induced by WSP. Transcription of canonical targets of the AHR, such as *CYP1A1* and *AHRR*, was robustly increased after just 30 minutes of WSP exposure, and we discovered novel AHR-regulated pathways and targets including the DNA methyltransferase, *DNMT3L*. Transcription of these genes and associated enhancers rapidly returned to near baseline by 120 minutes. The kinetics of AHR- and NFkB-regulated responses to WSP were distinguishable based on the timing of both transcriptional responses and chromatin remodeling, with induction of several cytokines implicated in maintaining NFkB responses through 120 minutes of exposure. In aggregate, our data establish a direct and primary role for AHR in mediating airway epithelial responses to WSP and identify crosstalk between AHR and NFkB signaling in controlling pro-inflammatory gene expression. This work also defines an integrated genomics-based strategy for deconvoluting multiplexed transcriptional responses to heterogeneous environmental exposures.

## Introduction

Particulate matter (PM) air pollution is estimated to cause over 4 million premature deaths annually on a worldwide basis (1). In addition to premature death, air pollution has been associated with a wide variety of health effects including acute and chronic respiratory symptoms, inflammation, exacerbations and morbidity (2–4). However, no specific therapies have been developed to address either acute or chronic health impacts of air pollution and our understanding of individual versus population risk from exposure is fragmentary. Compounding this, respirable particulate matter pollution is extremely heterogeneous (5).Thus, there is an important need to understand the effects of PM at the molecular level and to distinguish the cellular pathways that are regulated in response to different pollution exposures that are individually and in aggregate comprised of complex mixtures of chemical and particulates.

Respirable pollution is deposited along the airway and is known to exert a variety of effects on airway epithelial cells in a process that results in significant changes in gene expression (6,7).This has been investigated directly in human exposure studies (8), however, these processes are more frequently modeled using cell culture systems and standardized sources of pollution. For example, several pathways have been identified as regulating gene expression responses to lab-generated diesel exhaust particles (DEP) (9–11), including activation of the inflammasome, NFkB, and aryl hydrocarbon receptor (AHR) signaling (12–14). Inflammasome signaling, which results in NFkB activation, is associated with induction of cytokines and inflammatory responses(15). AHR, which is a bHLH-Pas domain transcription factor, is activated by a variety of potential AHR ligands that are byproducts of combustion (16). Upon ligand binding, AHR translocates to the nucleus and controls gene expression through binding to xenobiotic response elements (XREs) found in regulatory regions of canonical target genes (17–20), including detoxification enzymes such as CYP1A1, and the AHR repressor, AHRR. Functional XREs are found in regulatory regions for NFkB target genes, suggestive of a pro-inflammatory role in some contexts (21,22). However, AHR also induces the expression of IL24, a cytokine that represses NFkB(22). Thus, in addition to heterogeneity within air pollution, cellular responses to air pollution can encompass multiplexed transcriptional cascades whose crosstalk potentially results in both pro- and antiinflammatory effects.

One area of growing importance that further highlights challenges in contending with particle and chemical heterogeneity and cellular responses is wildfire pollution. As a consequence of climate changes, wildfires are increasing in size, prevalence, and impact on heavily populated areas (23,24). Wood smoke particles (WSP) derived from controlled combustion of wood have been used as a model to study cellular effects of wildfire smoke exposure (25–27). These WSP are composed of a heterogeneous mix of compounds including fine and coarse particulate matter, heavy metals, poly-aromatic hydrocarbons (PAHs) and volatile organic compounds (VOCs), which depend on the source of combustible material, similar to heterogeneity within bona fide wildfire pollution (26). Experimental WSP inhalation in healthy individuals recruits inflammatory cells, stimulates oxidative damage and impairs immune responses to viral infections. WSP exposure in cell culture models causes cytotoxicity, genotoxicity, cellular stress, formation of reactive oxygen species (ROS), and increased expression of pro-inflammatory cytokines (27–29). However, the molecular and transcriptional control of these effects has not been fully elucidated.

In this study, we developed a general approach to define the primary transcriptional mediators that result from exposing cultured human airway epithelial cells to a complex pollutant mix. We exposed Beas-2B airway epithelial cells to particles and compounds generated from controlled combustion of white oak. We assayed genome-wide nascent transcriptional responses and chromatin accessibility changes after 30 and 120 minutes of exposure to the white oak-derived WSP. We applied bioinformatics tools to cluster expression and epigenetic responses into distinct temporal patterns and to identify transcription factors that mediate these effects. We employed ChIP-qPCR, siRNA-mediated gene knockdown, and primary airway epithelial cells to validate transcriptional effectors discovered through this genomics-based discovery method.

## Methods

### Cell Culture

Beas-2B immortalized human bronchial epithelial cells were obtained from ATCC and cultured in Dulbecco’s Modified Eagle Medium (DMEM; Corning) with L-glutamine and 4.5 g/L glucose supplemented with 10% Fetal Bovine Serum (FBS; VWR) and 1% penicillin/streptomycin (Corning). De-identified primary human small airway epithelial cells (< 2 mm diameter; smAECs) were obtained from the National Jewish Health Biobank and plated onto an irradiated NIH/3T3 (ATCC) fibroblast feeder layer in F-medium containing 1 uM Y-27632 (APEX Bio). Upon visible colony formation (~7-10 days), smAECs were removed with 0.25% trypsin (Corning), plated on tissue culture dishes double-coated with Type I bovine collagen solution (Advanced Biomatrix), and grown to confluence in BronchiaLife Epithelial Medium supplemented with the complete LifeFactors Kit from Lifeline Cell Technology. All cells were maintained in 5% CO_2_ at 37°C.

### Wood Smoke Particle (WSP) Exposure

Wood Smoke Particles (WSP) were generated as describedand generously gifted by Dr. Andrew Ghio at the Environmental Protection Agency Laboratory in North Carolina (25). Briefly, wood smoke was generated by heating white oak wood on an electric heating element (Brinkmann Corporation, Dallas, TX) in a Quadra-fire 3100 woodstove (Colville, WA). The mean diameter of the freshly generated wood smoke particle was 0.14 micron. In the freshly generated particle, metal content was negligible. Particles were obtained by its mechanical disruption from the piping above the woodstove followed by sonication in water to disaggregate the particles. The ratio of elemental to organic carbon in the wood smoke particle was 0.004 (Sunset Laboratories, Hillsborough, NC) suggesting elemental carbon comprised approximately 0.4% of the particle. Diluted particle was injected into a gas chromatograph (Gerstel Thermal Desorption System/Agilent 6890 equipped with an Agilent 5973 mass selective detector). Identifiable compounds included levoglucosan (25.3%, mass/mass). After deposition and aggregation on the piping, there were measurable concentrations of metal including calcium (7.66 ppm), iron (0.76 ppm), magnesium (0.35 ppm), and aluminum (0.31 ppm) Particles were dehydrated for transportation and resuspended in PBS. Particles were disaggregated by sonication and sterilized under UV light for 20 minutes prior to addition directly to cell culture media at concentrations of .01 mg/ml to 1 mg/ml as indicated in the text.

### RNA purification and quantitative reverse transcription-PCR (qRT-PCR)

Beas-2B cells or smAECs were grown to confluence in 6-well tissue culture dishes (collagen-coated for smAECs, as described above) and treated with vehicle (PBS) or WSP for 2, 4 or 24 hours. Cells were harvested in TRIzol (Life Technologies) and RNA purified by PureLink RNA Mini Kit (Life Technologies) prior to qRT-PCR, performed with normalization to *RPL19* as previously detailed (30).Sequences of primers used for qRT-PCR analysis are provided in Supplemental Table S1.

### Precision Run-on Sequencing (PRO-seq)

Beas-2B cells were plated on 3 x 15 cm tissue culture dishes per treatment and grown to confluence, then treated with vehicle (PBS) or WSP for 30 or 120 minutes. Cells were harvested and nuclei prepared as reported previously (30). Aliquots of 10E6 nuclei were subjected to 3-minute nuclear run-on reactions in the presence of Biotin-11-CTP (PerkinElmer) and PRO-seq libraries were constructed in duplicate as described (31). Uniquely indexed libraries were pooled and sequenced on an Illumina NextSeq instrument using 75 bp single-end reads by the BioFrontiers Sequencing Facility at the University of Colorado Boulder.

### PRO-seq Computational Analysis

PRO-seq data were processed using a standardized Nextflow pipeline (https://github.com/Dowell-Lab/Nascent-Flow). A complete pipeline report detailing all software programs and versions utilized and a detailed quality control report including trimming, mapping, coverage, and complexity metrics are included in Supplemental File S1. TDF coverage files output by the pipeline, normalized by reads per million mapped, were visualized using the Integrative Genomics Viewer (IGV) genome browser (v. 2.8.0). FStitch (v. 1.0) and Tfit (v. 1.0) were used to identify regions with bidirectional transcriptional activity as described (30). Counts were calculated for each sorted BAM file using multiBamCov from the BEDTools suite (v. 2.28.0) and RefSeq:NCBI Reference Sequences for hg38 downloaded from the UCSC genome browser (May 18, 2018). Genes and lncRNAs were then filtered such that only the isoform with the highest number of reads per annotated length was kept, and DESeq2 (v. 1.20.0, Bioconductor release v. 3.7) was used to determine differentially transcribed genes between treatments. Gene clustering was then performed by intersecting differentially transcribed gene lists to form early-peak (upregulated in 30 min WSP vs vehicle and downregulated in 120 min vs 30 min WSP), early-plateau (upregulated in 30 min WSP vs vehicle and up in 120 min WSP vs vehicle) and late (upregulated in 120 min WSP vs vehicle and upregulated in 120 min vs 30 min WSP) temporal clusters. Functional annotation of differentially regulated genes was performed using DAVID v. 6.8 (32). For bidirectional comparisons, all predicted bidirectional Tfit calls were aggregated using mergeBed (argument-d 60) from BEDTools (v. 2.28.0) to generate an annotation file. Counts were then calculated for each sample using multicov (BEDTools v. 2.28.0), and DESeq2 was used to identify differentially transcribed bidirectionals between treatments.

### Assay for Transposase Accessible Chromatin using Sequencing (ATAC-seq)

Beas-2B cells were grown to confluence in 6-well tissue culture dishes and treated with vehicle (PBS) or WSP for 30 or 120 minutes. Cells were rinsed and scraped in ice-cold PBS, then ~50K cells from each treatment were pelleted and processed in duplicate for Omni-ATAC-seq using the protocol developed by Corces et al (33). Uniquely indexed libraries were pooled and sequenced on an Illumina NextSeq using 37 bp paired-end reads by the BioFrontiers Sequencing Facility at the University of Colorado Boulder.

### ATAC-seq Computational Analysis

ATAC-seq reads were trimmed for adapters, minimum length, and minimum quality using the bbduk tool from the BBMap Suite (v. 38.73) with arguments ‘ref=adapters.fa ktrim=r qtrim=10 k=23 mink=11 hdist=1 maq=10 minlen=20’. Quality control was monitored both pre- and post-trim for all samples using FastQC (v. 0.11.8). Trimmed reads were mapped to the human genome (hg38; downloaded from the UCSC genome browser on September 16, 2019, with corresponding hisat2 index files) using hisat2 (v. 2.1.0). Resulting SAM files were converted to sorted BAM files using samtools (v. 1.9) and to bedGraph coverage format using genomeCoverageBed from the BEDTools suite (v. 2.29.2). Read coverage was then normalized to reads per million mapped using a custom python script and files converted to TDF format using igvtools (v. 2.5.3) for visualization in IGV. MACS2 (v. 2.1.4) callpeak with ‘--SPMR’ argument was applied to sorted BAM files for each replicate pair of samples with parameters set to call peaks with log_2_ fold change > 1 above baseline with q < 1e-5 as significant. Peaks were then assigned to early-peak, early-plateau, and late temporal clusters based on simple presence/absence criteria within indicated treatment groups, illustrated by Venn diagrams in Figure 4.

**Figure 1.**
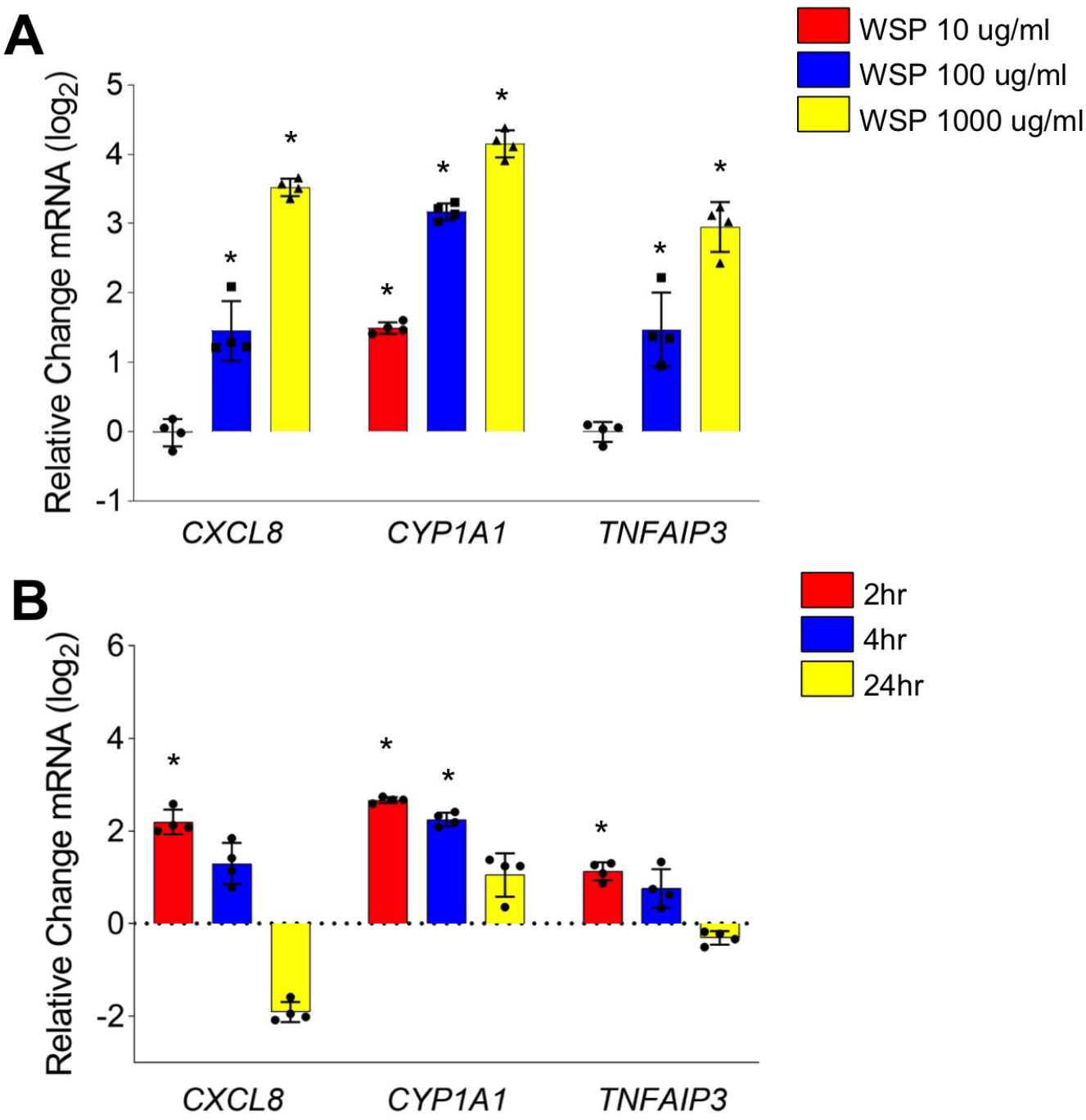
WSP exposure induces inflammatory gene transcription in Beas-2B airway epithelial cells. qRT-PCR analysis of indicated gene expression in Beas-2B cells treated with (A) WSP at indicated concentrations for 2 hours and (B) WSP at 100 ug/ml for 2, 4 and 24 hours. Bars represent mean normalized C_T_ values on a log_2_ scale (+SD) relative to vehicle-treated controls (n = 4/group, \**p* < 0.05 vs. vehicle).

**Figure 2.**
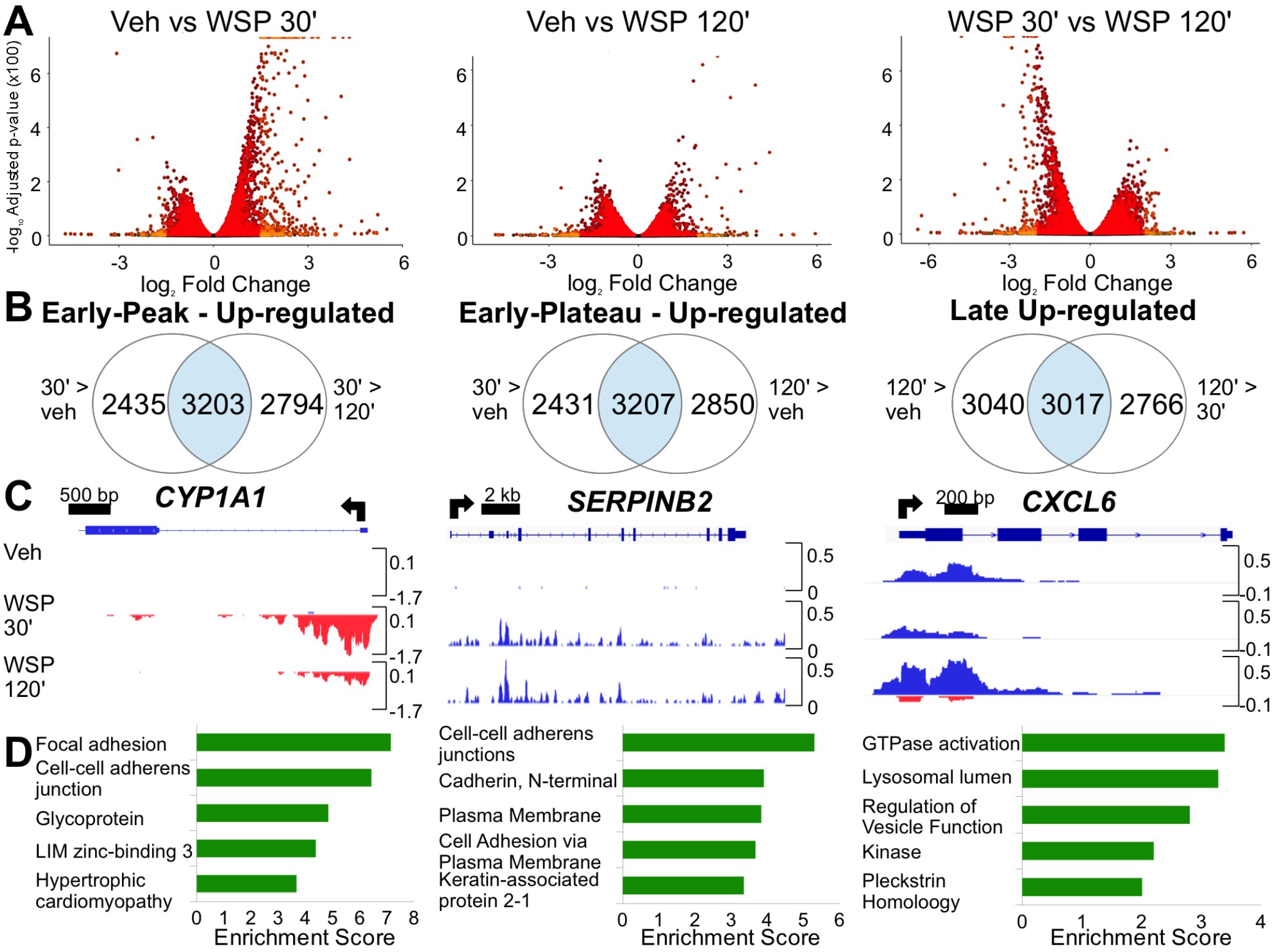
PRO-seq reveals distinct temporal patterns of rapid and transient transcriptional responses to WSP exposure. (A) Volcano plots illustrate differentially regulated nascent transcripts in Beas-2B cells treated with vehicle or WSP for 30 or 120 minutes and compared as indicated. Each point represents one gene, with light orange indicating p_adj_ < 0.05 and log_2_ fold change > 1, and red signifying differences not meeting both these criteria. Points justified to top of y-axis represent genes with an infinitesimally small p_adj_ that rounds to 0. (B) Criteria used to cluster genes by temporal kinetics of transcriptional induction by WSP and Venn diagrams showing number of differentially regulated transcripts (based on p_adj_ < 0.05) meeting these criteria. (C) Representative examples of each temporal cluster characterized in (B) shown as PRO-seq tracks visualized in the Integrative Genomics Viewer (IGV) genome browser based on counts per million mapped reads (vertical scales). Positive (blue) indicates reads annotated to the sense strand while negative (red) peaks reflect reads annotated to the antisense strand. The TSS and direction of transcription are indicated by arrows at the top of each panel. (D) Bar graphs display top 5 most significantly enriched functional annotation terms output by DAVID Functional Annotation Clustering applied to each set of WSP-induced transcripts characterized in (B).

**Figure 3.**
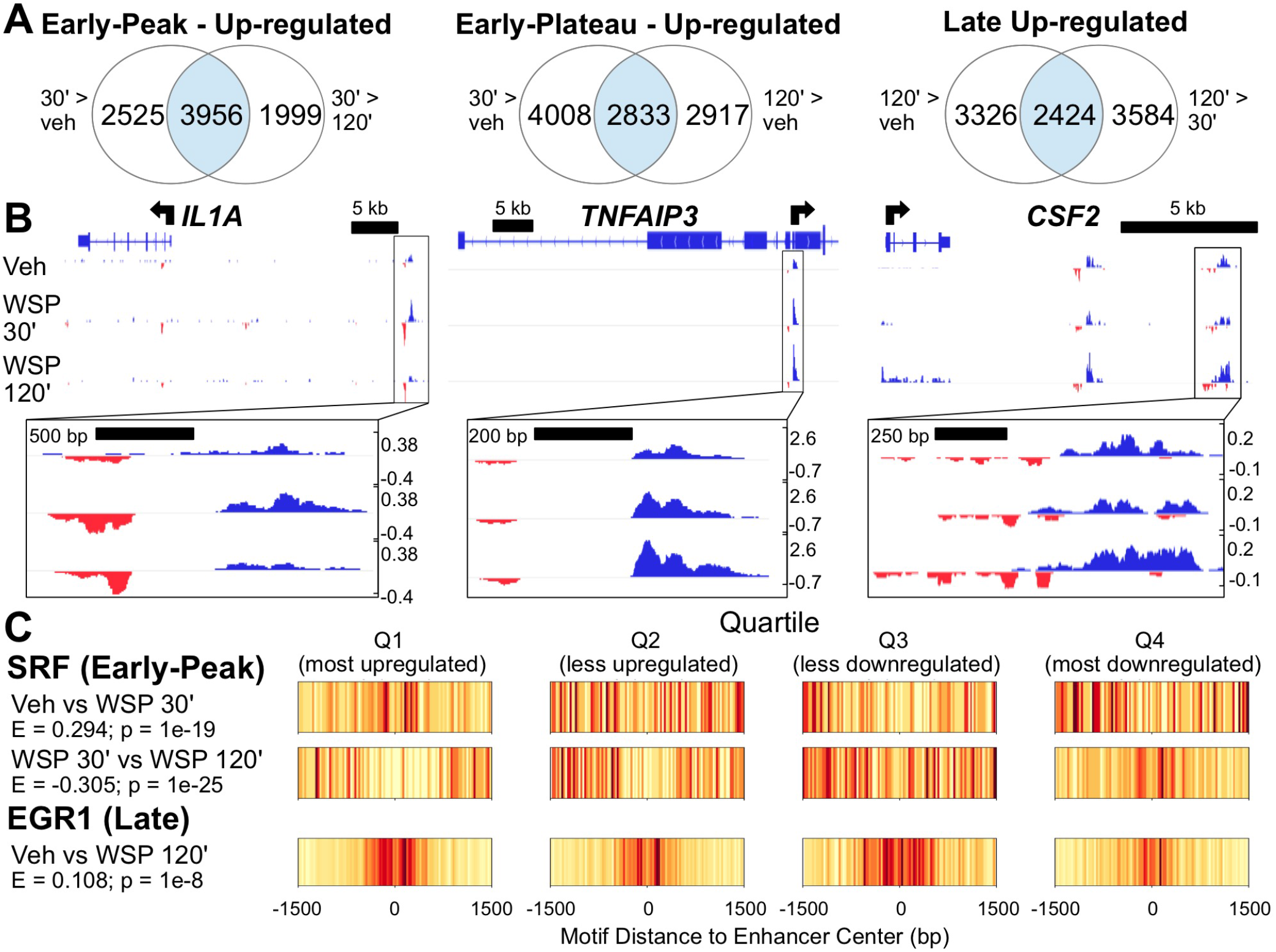
Genome-wide mapping of WSP-induced enhancer transcription defined by PRO-seq bidirectional signatures of RNA Polymerase II activity. (A) Venn diagrams indicating number of differentially regulated bidirectional transcripts (based on p_adj_ < 0.05) meeting indicated temporal clustering criteria, as described for Figure 2B. (B) IGV-visualized PRO-seq tracks of Tfit-called bidirectionals (boxed in black and magnified below each panel) and nearest genes exhibiting a similar pattern of regulation as representative examples of each temporal cluster characterized in (A). (C) Motif displacement distributions of significantly enriched transcription factor binding motifs within early-peak (top) versus late (bottom) clusters of differentially transcribed bidirectionals, as identified using TFEA. Each column represents the frequency of the indicated motif instance at the specified distance from the bidirectional center (labeled 0), where color specifies a normalized frequency relative to the entire 3 kb region. Darker colors indicate greater enrichment on a 0-1 scale.

**Figure 4.**
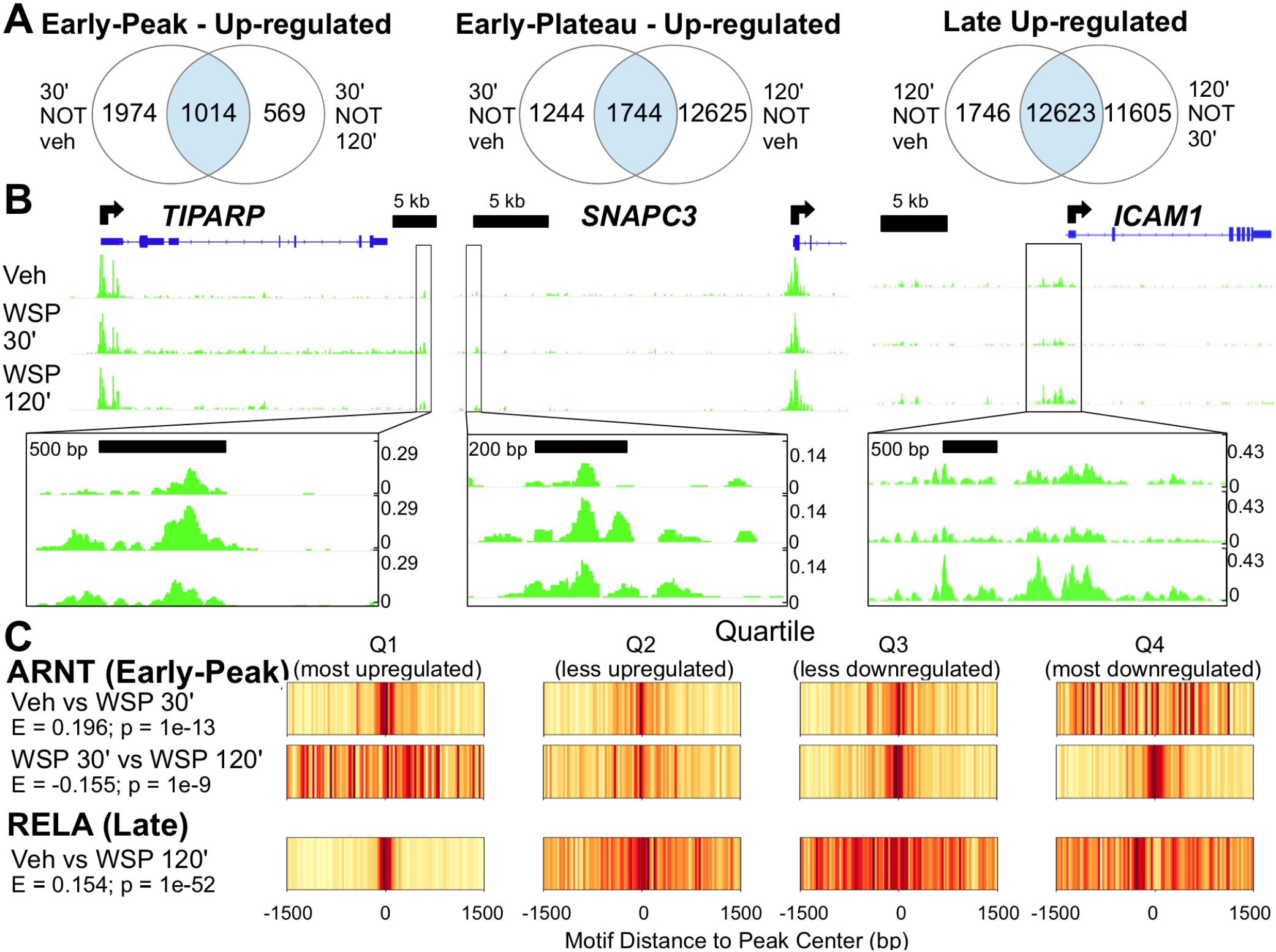
ATAC-seq demonstrates consistent temporal dynamics between chromatin accessibility and transcriptional responses to WSP and predicts key regulatory factors and pathways. (A) ATAC-seq was performed following exposure of Beas-2B cells to vehicle or WSP for 30 or 120 minutes. Venn diagrams indicate number of differentially regulated ATAC-seq peaks (based on p_adj_ < 0.05) meeting the indicated temporal clustering criteria, as described above. (B) IGV-visualized ATAC-seq tracks with MACS2-called peaks (boxed in black and magnified below each panel) and nearest genes exhibiting a similar pattern of regulation as representative examples of each temporal cluster characterized in (A). (C) Motif displacement distributions, as described for Figure 3C, from TFEA of differentially regulated MACS2-called ATAC-seq peaks representative of early-peak (top) and late (bottom) differential motif enrichment.

### Differential Transcription Factor Motif Enrichment

Transcription factor enrichment analysis for both PRO- and ATAC-seq datasets was performed using TFEA (v. 1.0; https://github.com/Dowell-Lab/TFEA; (34)). Sets of consensus regions of interest (ROIs) are first defined for all Tfit-called bidirectionals (for PRO-seq) or all MACS2-called peaks (for ATAC-seq) using the muMerge function of TFEA. TFEA then calculates read coverage for each replicate within each ROI and applies DESeq2 to determine differentially represented ROIs by treatment condition and assign rank as a function of p-value. For each ranked ROI, the FIMO tool (35)from the MEME suite scans the 3 kb sequence surrounding the ROI center (bidirectional origin or ATAC-seq peak summit) for positions of transcription factor consensus motif matches (with p-value cutoff of 10-5), represented by a curated set of position weight matrices (36)Enrichment scores (E-scores) for each motif are then calculated and corrected for sequence content to reduce known biases associated with local GC enrichment and p-values are determined using Z-scores. Motif displacement distributions (as in Figure 3C) are displayed as heatmaps where the location of all motif instances within +1500 bp are indicated relative to the ROI center. The intensity of color is a normalized fraction of motif instances within that bp range relative to the total number of motifs in the entire 3 kb region.

### Chromatin Immunoprecipitation (ChIP)-qPCR

Beas-2B cells were grown to confluence in 10 cm tissue culture dishes and treated with vehicle or WSP for 30 or 120 minutes. Cells were crosslinked by adding 16% methanol-free formaldehyde to a final concentration of 1% and incubating for 5 minutes at room temperature, then ChIP-qPCR was then performed as described (37) using 35 cycles of sonication. A mouse monoclonal antibody raised against AHR was purchased from Santa Cruz (sc-133088 X) and used at a concentration of 5 ug/sample. Assays were performed in biologic quadruplicate. ChIP-qPCR primer sequences are listed in Supplemental Table S2.

### Western Blotting

Western blotting and protein detection were performed using standard protocols (38)in unstimulated Beas-2B cells cultured to confluence in 6-well plates or following siRNA transfection, as detailed below. Primary antibodies purchased from Santa Cruz included anti-AHR (sc-133088X), anti-NFKB p65 (sc-372), and anti-GAPDH (sc-25778). Secondary antibodies were ECL Donkey anti-Rabbit IgG, HRP-linked F(ab’)_2_ fragment (NA9340) and ECL Sheep anti-Mouse IgG, HRP-Linked Whole Ab (NA931), both from GE Healthcare/Amersham.

### siRNA-mediated Gene Knockdown

Beas-2B cells were plated in 6-well tissue culture dishes in antibiotic-free medium. Approximately 24 hours later, cells were transfected with ON-TARGETplus siRNA SMARTpools (Horizon/Dharmacon) against human AHR (*si-AHR*; L-004990-00), RELA (*si-RELA*; L-003533-00), or a scrambled control (si-Ctrl; D-001810-10), at a final concentration of 25 nM using Lipofectamine RNAiMAX transfection reagent (Life Technologies) as instructed by the manufacturer. Media was replaced with fresh complete ~18 hours post-transfection, then 24 hours later, cells were treated with vehicle or WSP for 2 hours for qRT-PCR assays or harvested without additional stimulation for verification of knockdown via Western Blotting.

### Statistics

Statistical comparisons for qRT-PCR and ChIP-qPCR assays were made by 2-tailed t-test using the Bonferroni correction when appropriate. These analyses were conducted using statistical software embedded in Open Office (Apache OpenOffice v 4.1.7 http://www.openoffice.org/welcome/credits.html).

### Data Availability

Raw and processed sequencing data have been submitted to the NCBI Gene Expression Omnibus (GEO; https://www.ncbi.nlm.nih.gov/geo/) under accession GSE167372.

## Results

### WSP exposure causes rapid and temporally dynamic changes in gene transcription

As an initial step in developing a system to identify transcriptional mechanisms resulting from exposure to air pollution, we exposed Beas-2B cells, a transformed human bronchial epithelial cell line, to WSP in submerged culture at varying concentrations of 10, 100 and 1000 ug/ml (Figure 1A). WSP is an established model of wildfire pollution that is known to include significant chemical and particulate heterogeneity. We analyzed changes in gene expression after 2, 4 and 24 hours of exposure using qRT-PCR, focusing on previously identified targets of particulate matter in airway cells and NFkB target genes (38–41). We observed rapid induction of three candidate exposure-regulated genes (Figure 1B), with peak expression observed after just 2 hours of treatment. Thus, airway epithelial cells exhibit prompt, dynamic and dose-dependent changes in gene expression in response to WSP.

To explore the earliest transcriptional responses to WSP and determine underlying mechanisms, we conducted nascent transcript analysis of Beas-2B cell responses to WSP using PRO-seq (42). PRO-seq measures gene transcription with high temporal resolution based on RNA polymerase II activity and also allows for quantification of enhancer activity based on bidirectional signatures of enhancer RNA (eRNA) transcription (43,44). For these studies, we selected the highest tested dose of WSP (1000 ug/ml) in order to elicit maximal transcriptional responses for further bioinformatics analyses To define direct transcriptional responses to WSP and their initial dynamics, we analyzed nascent transcription after 30- and 120-minute exposure times relative to treatment with vehicle (PBS). We initially focused our analysis on changes in gene transcription in relationship to exposure time. As visualized in volcano plots (Figure 2A), at the 30-minute time point, increased transcription was the dominant response to WSP exposure, with many of these rapidly induced targets exhibiting reduced expression at 120 minutes relative to 30 minutes. We subsequently defined three distinct temporal clusters of transcriptional induction in response to WSP, as indicated by Venn diagrams in Figure 2B and detailed in the Methods, encompassing early-peak, early-plateau, and late clusters of differentially regulated gene transcripts. Complete gene lists for each cluster are presented in Supplemental Tables S3-S5, respectively. PRO-seq data were visualized in the Integrative Genomics Viewer (IGV) genome browser (45) and representative data tracks for each temporal cluster are shown in Figure 2B.

Next, to determine whether the different clusters represent distinct biologic processes regulated by WSP as a function of time, we performed functional annotation of the clusters using DAVID (32).Although there were annotation differences between all three clusters (Figure 2C; complete output for each cluster is listed in Supplemental Tables S6-S8), comparison of the late cluster to the early-peak and early-plateau annotations showed significant divergence in enriched pathways. In addition, annotation of the early-peak cluster identified enrichment of AHR-related pathways including Cytochrome P450 metabolism (p = 6.438e-5) and PAS domaincontaining transcription factors (p = 3.399e-5). Functional annotation of early-plateau cluster genes also identified enrichment of PAS domain-containing transcription factors (p = 7.424e-4) and NFkB signaling pathways (p = 9.775e-3). In aggregate, our analysis of nascent transcription indicates that WSP exposure causes a rapid remodeling of airway epithelial gene expression and associated cellular processes. These changes are temporally dynamic, and ontology analysis implicates AHR and NFkB as potential transcriptional mediators of early effects of WSP exposure on transcription.

### Enhancer RNA (eRNA) transcription is regulated by WSP

To further study mechanisms underlying regulation of gene expression by WSP, we next analyzed changes in enhancer expression by utilizing Tfit to identify bidirectional signatures of RNA polymerase II activity associated with nascent eRNA and promoter transcription (46). We then used DEseq2 to determine differential expression of these bidirectional transcripts following vehicle or WSP exposure at both time points. Through this analysis we defined clusters of bidirectional transcriptional regulation in response to WSP analogous to the clusters observed for gene transcription (Figure 3A). Examples of these distinct bidirectional transcript response patterns to WSP as visualized in IGV are shown in Figure 3B. Complete lists of differentially transcribed bidirectionals from early-peak, early-plateau, and late temporal clusters, as well as genomic coordinates for all identified bidirectional transcripts, are provided in Supplemental Tables S9-S12, respectively. Taken together, these data indicate that dynamic changes in airway epithelial cell gene expression in response to WSP are associated with rapid changes in enhancer activity.

Dynamic changes in enhancer activity generally result from differential transcription factor binding within the enhancer. Therefore, to identify transcription factors that control the primary response to WSP exposure, we interrogated enhancer regions whose activity, based on PRO-seq signatures, changed dynamically in response to WSP exposure. To accomplish this, we identified the origin of bidirectional transcription and applied the Transcription Factor Enrichment Analysis (TFEA) tool to these regions to identify differential motif enrichment for specific transcription factors (34). TFEA quantifies the degree of co-localization of transcription factor motif instances within the center of specific genomic regions, which in this case are sites of bidirectional transcription. We therefore compared changes in enrichment (E-) scores between vehicle and 30 minutes, 30 minutes and 120 minutes, and vehicle and 120 minutes, analogous to the three temporal clusters we previously defined based on induction of transcription. Motifs for the TCF-SRF complex, which is activated in response to ERK and MAPK signaling (47,48), were identified as significantly enriched in the 30-minute dataset in comparison to vehicle (e.g. Fig 3C, top), suggesting a rapid kinase-mediated transcriptional response to WSP. Comparison of 30 minutes to 120 minutes showed that enrichment of TCF and SRF motifs returned to baseline by 120 minutes, whereas EGR family motifs were enriched at 120 minutes in comparison to vehicle (Figure 3C, bottom). These data suggest that dynamic changes in transcription factor utilization underlie different kinetics in transcriptional responses to WSP. Significantly enriched and depleted motifs for each treatment comparison are summarized in Supplemental Table S13.

### Changes in chromatin structure identify AHR as driving early transcriptional responses to WSP

Some transcription factors are more associated with chromatin remodeling than directly altering RNA polymerase activity, and TFEA can also be applied to regions based on changes in chromatin structure to identify these regulatory pathways (34). Therefore, to determine effects of WSP exposure on chromatin status, we performed the Assay for Transposase Accessible Chromatin using sequencing (ATAC-seq) in Beas-2B cells following 30 and 120 minutes of WSP exposure (33,49).Based on MACS2-defined peaks (50), we partitioned the ATAC-seq data temporally into the time-dependent clusters we had previously defined for nascent transcription (Figure 4A), noting that the number of new peaks occurring after 120 minutes of WSP was significantly greater than the number of new peaks at 30 minutes. Examples of loci that fall into each cluster are shown in Figure 4B. These data indicate that WSP causes significant chromatin remodeling, and the extent of chromatin remodeling in response to WSP increases over time.

As a further approach to define transcriptional mediators of genomic responses to WSP, we interrogated 3 kb regions centered on MACS2-defined ATAC-seq peaks using TFEA (51)to compare motif enrichment based on treatment and time. This analysis identified strong enrichment for the ARNT motif, the obligate AHR dimerization partner. This enrichment was evident at 30 minutes but decreased to baseline by 120 minutes (Figure 4C, top). In contrast, differential motif enrichment for RELA, a member of the NFkB complex, was significantly increased at 120 minutes but not 30 minutes (Figure 4C, bottom). These data strongly implicate AHR signaling as mediating early transcriptional responses to WSP in association with rapid chromatin remodeling, whereas effects of NFkB appear to peak at a later time.

### Novel targets of AHR signaling defined through integrated genomics

In light of the known association of other forms of PM pollution with AHR signaling (12), and the identification of AHR through applying unbiased bioinformatics approaches to our genome-wide data, we scrutinized the early-peak cluster for canonical AHR target genes. We found a significant increase in transcription of numerous AHR targets at 30 minutes of WSP exposure that sharply decreased after 120 minutes (Table 1). Examples of integrated PRO-seq and ATAC-seq data for several established AHR targets, such as *CYP1B1*, and their associated enhancers visualized in IGV are shown in Figure 5A. MatInspector analysis of each of these dynamically regulated enhancer regions revealed matches for the canonical AHR/ARNT response element (52), implicating AHR as likely mediating these rapid transcriptional effects. We then identified additional genes of interest demonstrating early peak kinetics (see Table 1) and interrogated these genes for canonical AHR binding sites in regulatory regions. *DNMTL3*, which is reported to function within a protein complex to induce DNA methylation (53), is amongst the putative novel AHR targets we identified in this manner.

**Figure 5.**
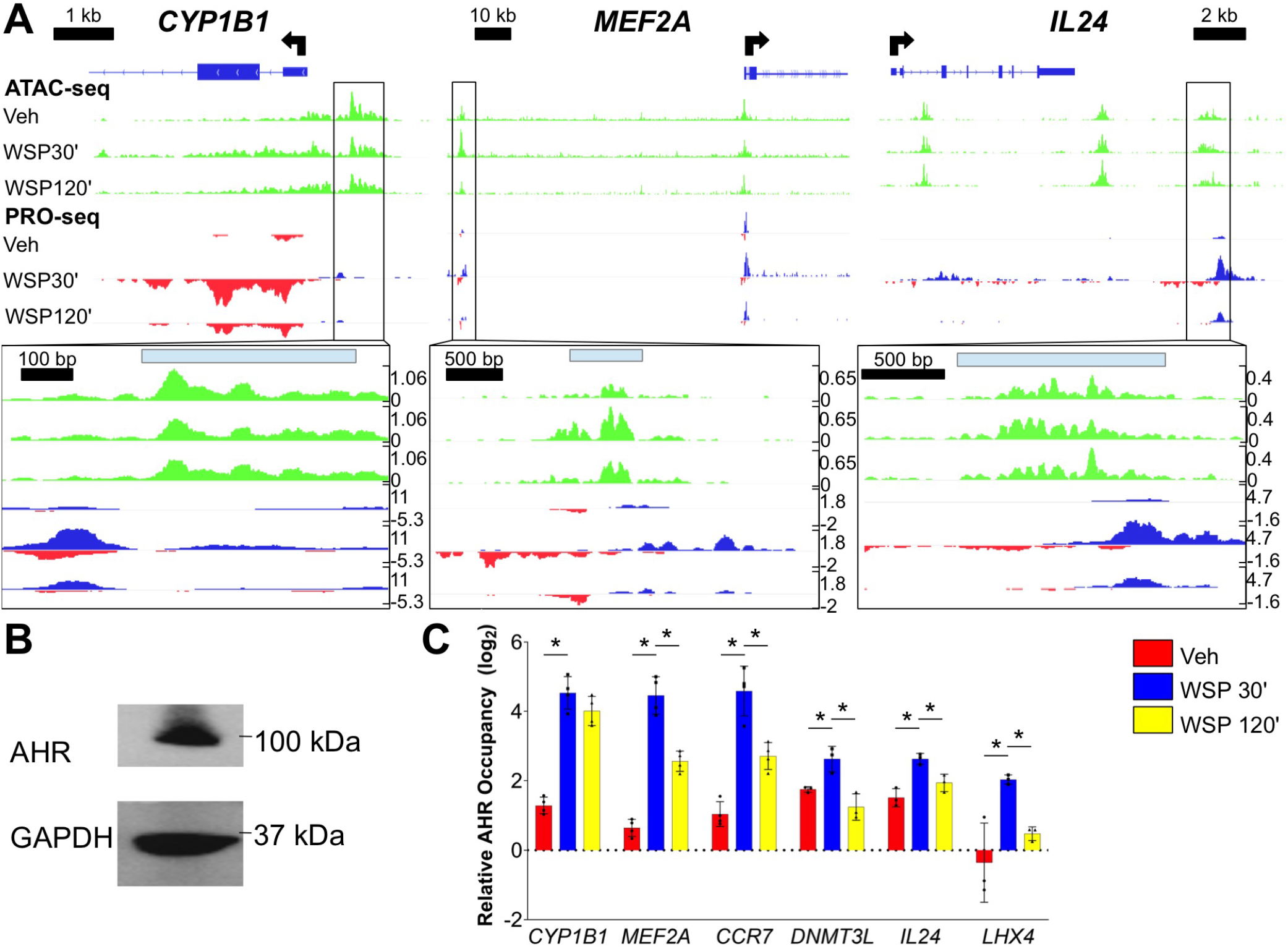
Integrated analysis of chromatin remodeling and nascent eRNA profiling datasets identifies sites of direct AHR occupancy and functional regulation of early-peak transcriptional targets of WSP. (A) ATAC-seq and PRO-seq tracks aligned in IGV at early-peak WSP response genes that are known targets of AHR signaling. Associated early-peak eRNAs and ATAC-seq peaks are boxed and magnified below each panel, with locations of matches to the consensus AHR binding motif, determined using MatInspector, indicated by light blue bars. (B) Western blot verification of AHR protein expression in Beas-2B cells under basal culture conditions; GAPDH was a loading control. (C) qPCR primers were designed for the three regions examined in (A) plus one additional canonical (*CCR7*) and two novel (*DNMT3L*, *LHX4*) early-peak targets of AHR, then ChIP-qPCR was performed in Beas-2B cells following exposure to vehicle or WSP for 30 or 120 minutes. Bars represent AHR occupancy on a log_2_ scale (+SD), expressed as the mean C_T_ value at each target region relative to the geometric mean of C_T_ values at three negative control regions (n = 4/group, \**p* < 0.05 for indicated comparison).

**Table 1:** Selected targets from early-peak cluster of WSP-induced genes. P_a_dj values listed as 0 indicate infinitesimally small p_adj_ values that round to 0.

|  | Vehicle vs WSP 30 min |  | WSP 30 min vs WSP 120 min |  |
| --- | --- | --- | --- | --- |
| Gene | Log<sub>2</sub> Fold Change | $p_{adj}$ | Log<sub>2</sub> Fold Change | $p_{adj}$ |
| *IL24* | 5.757 | 0 | -2.359 | 6.81E-141 |
| *EGR1* | 5.491 | 0 | -4.790 | 0 |
| *KRT17* | 5.128 | 0 | -2.467 | 0 |
| *LHX4* | 5.066 | 0 | -2.421 | 3.72E-152 |
| *MAFF* | 4.412 | 0 | -2.869 | 0 |
| *TAGLN* | 4.169 | 0 | -2.988 | 0 |
| *CCN2* | 3.798 | 0 | -3.018 | 0 |
| *SPOCD1* | 3.629 | 0 | -1.928 | 1.75E-158 |
| *TIPARP* | 2.777 | 0 | -1.975 | 1.37E-282 |
| *SRF* | 2.773 | 0 | -2.202 | 5.42E-218 |
| *MYH9* | 2.756 | 0 | -1.797 | 6.60E-282 |
| *CYP1B1* | 2.508 | 0 | -1.384 | 9.16E-126 |
| *DNMT3L* | 3.802 | 8.63E-280 | -2.116 | 1.34E-116 |
| *AHRR* | 1.794 | 2.22E-238 | -0.737 | 4.55E-41 |
| *CYP1A1* | 5.410 | 1.51E-224 | -1.466 | 4.35E-45 |
| *ALDH3A1* | 2.512 | 2.58E-124 | -0.992 | 1.21E-22 |
| *ARNT2* | 1.262 | 5.65E-72 | -0.590 | 7.75E-17 |
| *MT2A* | 1.318 | 1.63E-64 | -1.328 | 1.34E-64 |
| *MEF2A* | 0.836 | 4.16E-35 | -0.994 | 6.95E-49 |
| *CCR7* | 2.153 | 1.43E-32 | -1.560 | 1.24E-18 |
| *CYP1A2* | 4.662 | 7.97E-11 | -2.324 | 9.82E-06 |

To definitively establish that AHR directs rapid and transient effects of WSP on airway epithelial gene expression, we performed ChIP-qPCR assays for AHR occupancy at several putative target loci. First, we confirmed by Western blot that AHR is expressed in this cell type (Figure 5B). Then we performed AHR ChIP-qPCR in Beas-2B cells following WSP exposure for 30 and 120 min. This demonstrated a gain in AHR occupancy in regulatory regions for canonical targets such as *CYP1B1*, as well as several novel targets, including *DNMT3L*, following exposure to WSP for 30 minutes (Figure 5C). AHR occupancy was decreased relative to the 30-minute time point after 120 minutes, consistent with the enhancer transcriptional signatures observed in the PRO-seq datasets (Figures 5A). Thus, AHR directly and rapidly activates gene expression in response to WSP, however, AHR occupancy and direct effects on gene expression peak before 2 hours.

### TCDD exposure confirms novel AHR targets in Beas-2B and primary airway epithelial cells

WSP contains a range of potential AHR ligands and whether the rapid AHR signaling response observed at observed at 1 mg/ml of WSP is relevant to established AHR agonists is not clear. Therefore, we exposed Beas-2B cells to 2,3,7,8-tetrachlorodibenzo-p-dioxin (TCDD), a prototypical ligand for AHR (20). We compared AHR induction between two doses of WSP and a standard 10 nM TCDD concentration (54). This demonstrated that expression of several, but not all early-peak target genes, is dependent on AHR activation and occurs in response to standard doses of TCDD and WSP (Figure 6A). To determine whether this also occurs in primary cells, we exposed primary small airway epithelial cells (Sm18) to TCDD and to WSP. These primary cells manifested similar expression responses to these stimuli, including with respect to novel AHR target genes (Figure 6B). AHR occupancy at regulatory regions neighboring several early-peak AHR target genes was also increased with exposure to standard doses of TCDD. Our findings thus establish a physiologically diverse set of AHR targets in response to both combustion-derived AHR ligands, with targets implicated in regulating DNA methylation (*DNMT3L*), inflammation (*IL24*), cell-cell adhesion (*LHX4*), glucocorticoid responses (*NR3C1*) and transcriptional regulation (*MAFF*).

**Figure 6.**
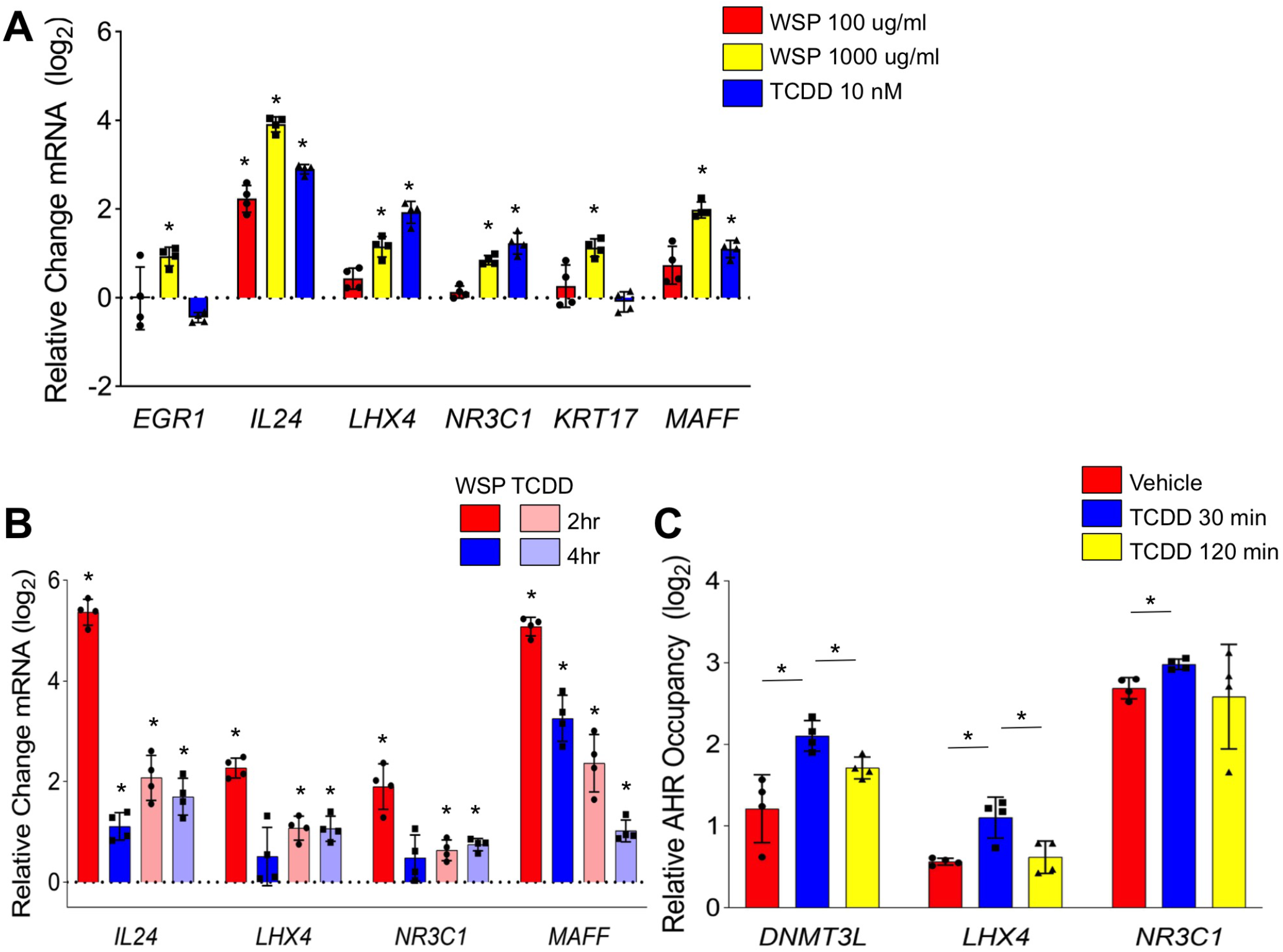
Exposure to TCDD induces transcription of AHR-dependent WSP targets in Beas-2B and primary airway epithelial cells. qRT-PCR analysis of indicated gene expression in (A) Beas-2B cells treated with two concentrations of WSP or TCDD (10 nM) for 2 hours and (B) primary human small airway epithelial cells treated with WSP (1000 ug/ml) or TCDD (10 nM) for 2 or 4 hours. Bars represent mean normalized CT values on a log_2_ scale (+SD) relative to vehicle-treated controls (n = 4/group, \**p* < 0.05 vs. vehicle). (C) ChIP-qPCR was performed in Beas-2B cells for novel early-peak targets of AHR following stimulation with TCDD (10 nM). Bars represent AHR occupancy on a log_2_ scale (+SD), expressed as the mean C_T_ value at each target region relative to the geometric mean of C_T_ values at three negative control regions (n = 4/group, \**p* < 0.05 for indicated comparison).

### Crosstalk between AHR and NFkB signaling in response to WSP

Our data, in accordance with literature on other sources of PM pollution, suggest that both AHR and NFkB respond to WSP, but whether there is direct crosstalk between these pathways is unclear. To study the role of AHR in inflammatory transcriptional responses to WSP, we analyzed gene expression responses to WSP in the setting of siRNA-mediated knockdown of AHR or the RELA subunit of NFkB. Using qRT-PCR, we quantified the expression of several canonical AHR targets, as well as several established targets of NFkB (Figure 7A). After WSP exposure for 2 hours, *AHR* knockdown cells (verified by Western blotting, Figure 7B) demonstrated decreased expression of *AHRR* and *CYP1A1* in comparison to cells treated with si-Ctrl. There was no effect of AHR knockdown on expression of *CXCL8*, a prototypical NFkB target. Surprisingly, WSP-mediated increases in *IL1A* and *IL1B* expression were attenuated with *AHR* knockdown, indicating a direct or secondary role for AHR in inducing these cytokines in response to WSP. Knockdown of *RELA* resulted in an essentially reciprocal pattern, with a complete abrogation of *CXCL8* induction in response to WSP, a minimal effect on the induction of *AHRR* and *CYP1A1*, and an intermediate effect on *IL1A* and *IL1B* (Figures 7C-D). Thus, AHR and NFkB are both required for maximal increases in *IL1A* and *IL1B* expression in response to WSP, indicating functional crosstalk between AHR and inflammatory responses to WSP. Moreover, as *IL1A* and *IL1B* induce NFkB activity, these data indicate convergence of AHR and inflammatory signaling in a positive feedback circuit that may promote further inflammation.

**Figure 7.**
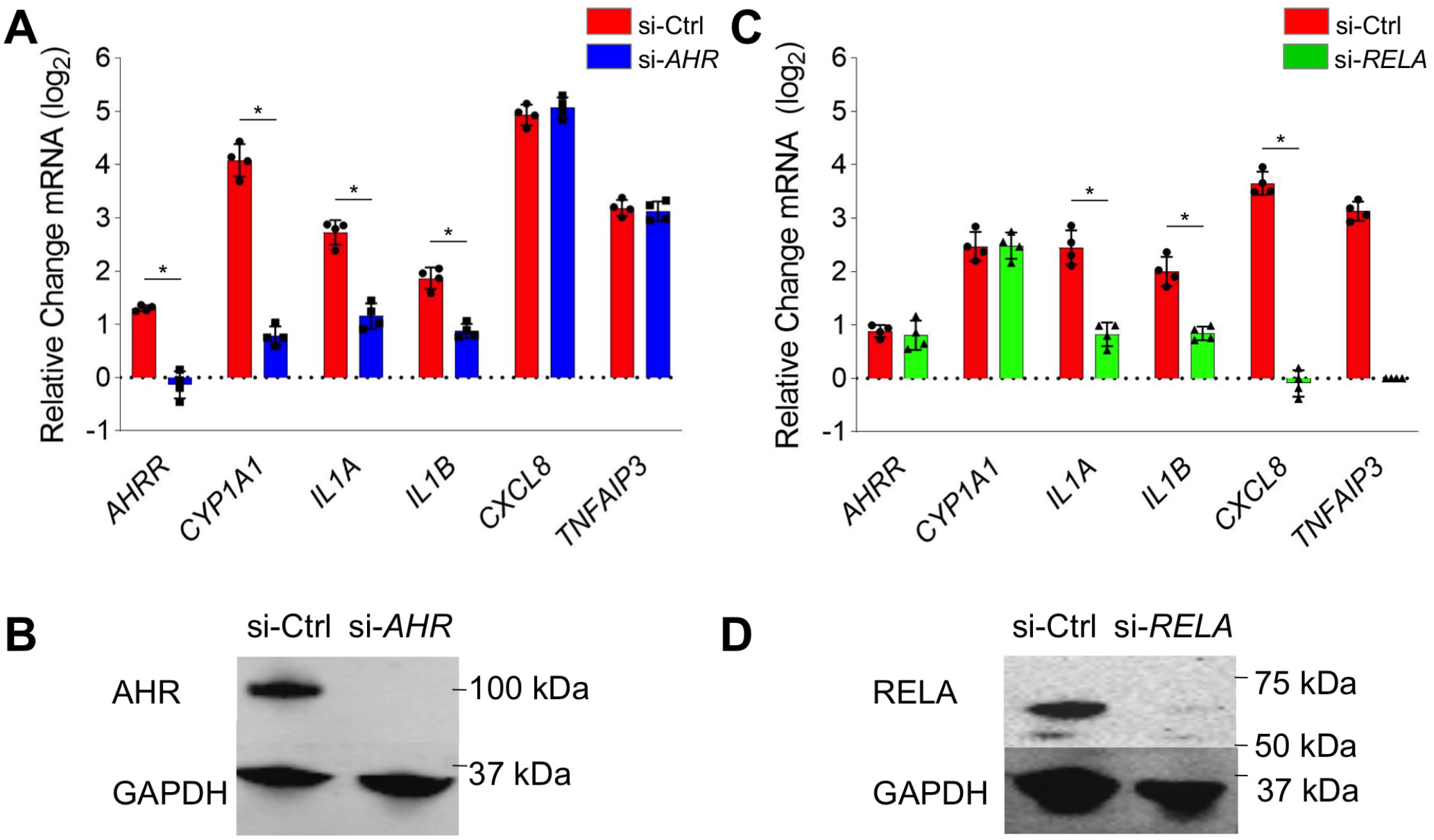
Airway epithelial inflammatory responses to WSP are mediated in part through complex crosstalk between AHR and NFKB. (A) Beas-2B cells transiently transfected with siRNA targeting AHR (si-*AHR*) or a scrambled control construct (si-Ctrl) were treated with vehicle or WSP for 2 hours and then assayed for indicated gene expression using qRT-PCR. Bars represent mean normalized C_T_ values on a log_2_ scale (+SD) relative to si-Ctrl+vehicle-treated controls (n = 4/group, \**p* < 0.05 for indicated comparison). (B) Western blot verification of AHR knockdown by si-*AHR* transfection. (C) si-*RELA*- or si-Ctrl-transfected Beas-2B cells were treated and assayed as described for (A). (D) Western blot verification of RELA knockdown by si-*RELA*.

## Discussion

Transcriptional regulation of airway epithelial responses to air pollution are challenging to define due to heterogeneity of the constituent respirable particulates and compounds. Through analyzing nascent transcription and performing ATAC-seq, here we show that airway epithelial cells undergo temporally dynamic changes in gene expression and chromatin structure in response to WSP, which models both wildfire smoke and exposure-related heterogeneity. Using unbiased bioinformatic approaches, we identified distinct roles for three transcription factor families, TCF/SRF, AHR, and NFkB, in mediating responses to WSP. We subsequently defined a set of canonical AHR targets that exhibit increased transcription after 30 minutes of WSP exposure in association with increased activity of nearby enhancers. Using ChIP, we established AHR occupancy at canonical XREs within enhancers and promoters for several target genes, and we further confirmed a role for AHR in mediating gene expression responses through knockdown experiments and exposure to TCDD, a well-established AHR ligand. In aggregate, our data establish a framework for identifying transcription factors that respond to air pollution and define a direct role for AHR in inducing rapid, transient transcriptional effects in response to WSP.

PAHs found within various forms of air pollution are known to function as AHR ligands (16), however, direct induction of AHR by wood smoke particulates had not been previously reported. By virtue of the high resolution afforded by integrating PRO-seq and ATAC-seq data, we identified canonical AHR target genes that are rapidly induced in Beas-2B cells following WSP exposure, thereby expanding our understanding of AHR signaling with respect to air pollution in airway epithelial cells. Rapidly induced targets included *CYP1A1* and *CYP1A2*, which metabolize AHR-activating PAHs that are found in WSP; the AHR repressor, *AHRR*; and *TIPARP*, an ADP-ribosyltransferase that represses AHR activity though ribosylation (55,56). Thus, in airway epithelial cells, at least three distinct negative feedback mechanisms regulated by AHR are implicated in the rapid decrease in transcription of primary AHR targets following WSP exposure. We also defined a novel set of genes regulated by AHR based on several criteria including: 1) rapid, transient transcriptional induction patterns similar to that observed for canonical AHR targets with WSP exposure, 2) the discovery of WSP-responsive enhancers for these genes harboring centrally located canonical XREs, and 3) confirmation of target gene activation following TCDD exposure.

In contrast to the powerful negative feedback system that limits the duration of direct effects of AHR on gene transcription, a number of genes we identified are candidates to mediate prolonged secondary responses to WSP-mediated activation of AHR signaling. For example, transcription of the MEF2 family of transcription factors was induced after 30 minutes of WSP exposure. Based on TFEA indicating enrichment for motifs for each of these factors among active enhancers at 120 minutes versus vehicle (FDR < 0.05), our data implicate the MEF2 family in mediating secondary gene expression responses to WSP. In addition to this family, the bZip transcription factor, *MAFF*, *LHX4*, a lim homeobox factor, and the cytokine, *IL24*, which is implicated in lung and airway remodeling in response to different inflammatory stimuli, were also amongst the set of AHR targets we identified (57–60).Thus, the short-lived primary transcriptional response to WSP mediated by AHR includes targets that likely contribute to more sustained changes in gene expression, possibly in collaboration with other rapid transcriptional effectors of the response to WSP, such as the TCF/SRF family. Although we observed induction of AHR targets in primary human small airway epithelial cells and similar kinetics, future work is needed to more fully establish the relevance and temporal properties of AHR signaling responses to WSP in primary, differentiated airway epithelial cells cultured at air-liquid interface. How AHR signaling in response to a heterogenous exposure that results in the multiplexed transcriptional cascade activity characterized here compares to induction of AHR signaling in response to a single ligand, such as TCDD, also remains to be determined.

Persistent epigenetic changes such as DNA methylation, which can exert a memory effect on cell phenotype through modifying gene expression, are believed to contribute to long-term health effects of exposure to pollutants (61,62). Through PRO-seq, we identified rapid and transient induction in response to WSP of the DNA methylase, *DNMTL3* (*63*) Although PAH-containing pollutants have been previously reported to cause increased DNA methylation (64,65), we are unaware of any reports that have identified specific methylation pathways that might mediate this epigenetic response to WSP or other pollutants containing AHR ligands. Whether this pathway is responsible for teratogenic and other long-term effects of AHR signaling, including altered DNA methylation, remains to be confirmed in future work.

In addition to AHR, we also identified NFkB as a mediator of transcriptional and chromatin responses to WSP in airway epithelial cells. Canonical targets of NFkB that were activated in response to WSP include *TNFAIP3*, *CXCL8*, and *IL1B*, among others. Knockdown of *RELA* confirmed that induction of *TNFAIP3* and *CXCL8* was largely dependent on the NFkB complex. Although other studies have implicated inflammasome activation in response to particulates as directly inducing NFkB activity (66), the mechanisms underlying NFkB activation in response to WSP remain to be elucidated. The timing of AHR and NFkB transcriptional motif utilization based on TFEA, however, suggests that NFkB-mediated responses to WSP occur in a temporally distinctive process from AHR signaling. Specifically, we observed significantly increased central NFkB motif enrichment within ATAC-seq peaks at the 120-minute time point relative to 30 minutes, whereas AHR motif utilization peaked at 30 minutes. Given the rapid induction of *IL1B* and *CXCL8* in response to WSP, both of which are known activators of NFkB signaling, NFkB activity in response to WSP exposure is likely partially attributable to positive feedback mediated through these and other cytokines that activate NFkB.

The identification of novel AHR targets and AHR crosstalk with NFkB are two important aspects of the transcriptional response to WSP that we captured in this study. Our approach also comprehensively mapped the diverse gene targets and transcriptional regulators that respond to WSP over two time points, including defining rapid and transient transcriptional responses mediated by several families of transcription factors in response to this exposure. We employed a WSP exposure in this study both to model wildfire particulates and to represent the multiplexed transcriptional programming that arises in consequence to respirable particle and compound heterogeneity, which is a defining characteristic of many forms of air pollution. Our study thus describes a novel and generalizable approach to partition heterogeneous pollutant exposures into discrete transcriptional response pathways (Figure 8). The strength of this method is supported by validation experiments we performed using conventional assays, such as qRT- and ChIP-PCR, and a single chemical ligand, TCDD, to activate AHR directly. Although further experiments are necessary to investigate the pathways identified by our approach and their physiologic consequences, in aggregate our data both define novel pathways that respond to WSP and inform a general strategy to deconvolute multiplexed transcriptional responses that arise from air pollution exposure.

**Figure 8.**
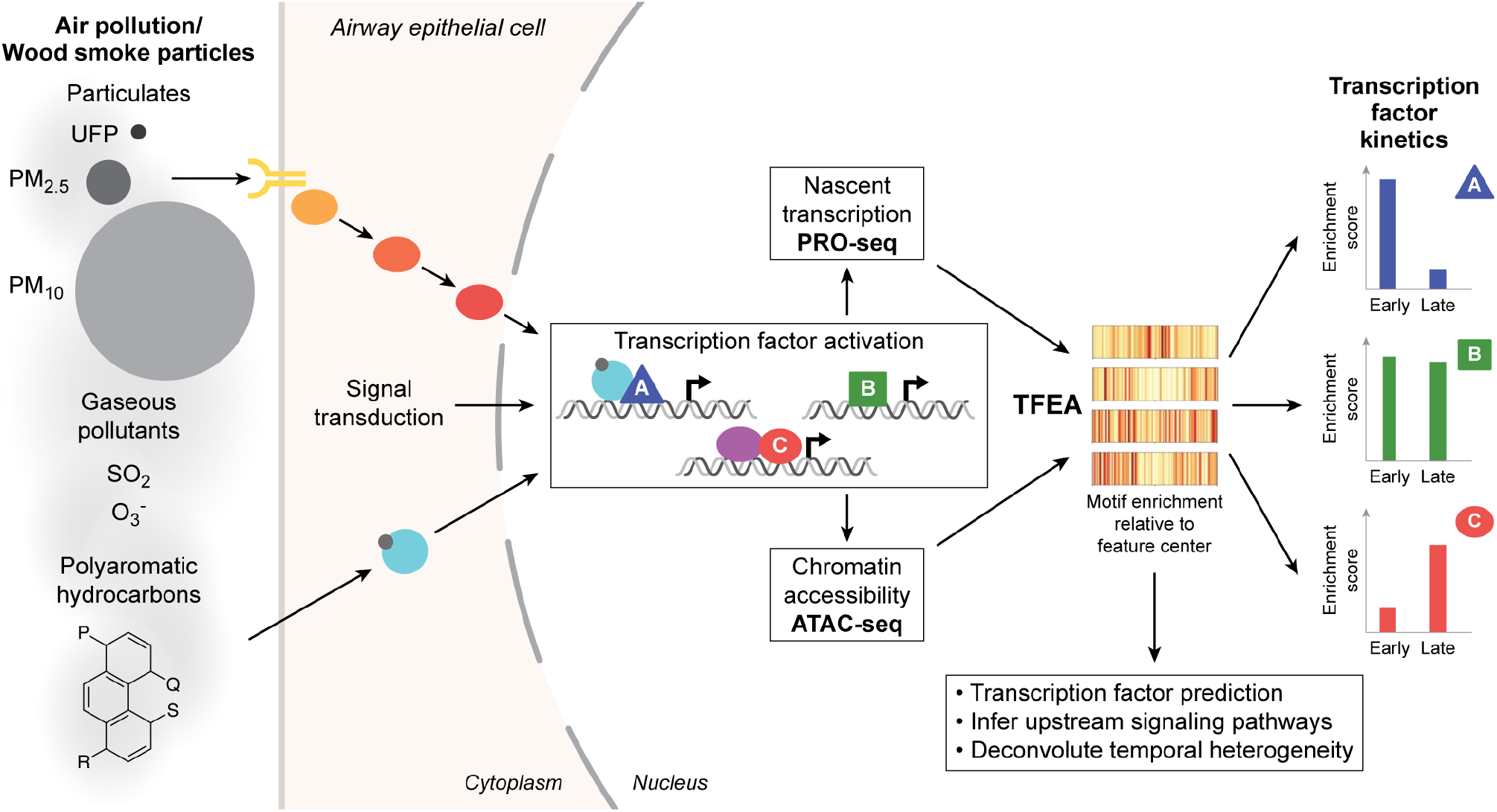
Genomics-based deconvolution of multiplexed transcriptional responses to complex air pollutants. WSP exposure is predicted to result in complex signal transduction events that lead to activation of multiple transcription factor pathways. To deconstruct these molecular processes, we generated paired genomewide chromatin accessibility and nascent transcription profiles in response to two WSP exposure time points. Unbiased transcription factor enrichment analysis (TFEA) of these data identified distinct families of transcription factors that mediate changes in gene expression, enhancer activity, and chromatin structure in response to WSP. Transcription factor activity could be further clustered based on timing, and in some cases, responses were rapid and transient. This system can be applied to identify and compare epigenetic and transcription factor responses to different sources of air pollution.

## Acknowledgments

The CU Boulder BioFrontiers Sequencing facility provided invaluable technical assistance.

## References

1. Cohen, A. J., Brauer, M., Burnett, R., Anderson, H. R., Frostad, J., Estep, K., Balakrishnan, K., Brunekreef, B., Dandona, L., Dandona, R., Feigin, V., Freedman, G., Hubbell, B., Jobling, A., Kan, H., Knibbs, L., Liu, Y., Martin, R., Morawska, L., Pope, C. A., 3rd, Shin, H., Straif, K., Shaddick, G., Thomas, M., van Dingenen, R., van Donkelaar, A., Vos, T., Murray, C. J. L., and Forouzanfar, M. H. (2017) Estimates and 25-year trends of the global burden of disease attributable to ambient air pollution: an analysis of data from the Global Burden of Diseases Study 2015. Lancet (London, England) 389, 1907–1918

2. Reid, C. E., Brauer, M., Johnston, F. H., Jerrett, M., Balmes, J. R., and Elliott, C. T. (2016) Critical Review of Health Impacts of Wildfire Smoke Exposure. Environmental health perspectives 124, 1334–1343

3. Adetona, O., Reinhardt, T. E., Domitrovich, J., Broyles, G., Adetona, A. M., Kleinman, M. T., Ottmar, R. D., and Naeher, L. P. (2016) Review of the health effects of wildland fire smoke on wildland firefighters and the public. Inhal Toxicol 28, 95–139

4. Wu, C. M., Adetona, A., Song, C., Naeher, L., and Adetona, O. (2020) Measuring acute pulmonary responses to occupational wildland fire smoke exposure using exhaled breath condensate. Archives of Environmental and Occupational Health 75, 65–69

5. Hime, N. J., Marks, G. B., and Cowie, C. T. (2018) A Comparison of the Health Effects of Ambient Particulate Matter Air Pollution from Five Emission Sources. International journal of environmental research and public health 15

6. O’Beirne, S. L., Shenoy, S. A., Salit, J., Strulovici-Barel, Y., Kaner, R. J., Visvanathan, S., Fine, J. S., Mezey, J. G., and Crystal, R. G. (2018) Ambient Pollution Related Reprogramming of the Human Small Airway Epithelial Transcriptome. Am J Respir Crit Care Med

7. Holloway, J. W., Savarimuthu Francis, S., Fong, K. M., and Yang, I. A. (2012) Genomics and the respiratory effects of air pollution exposure. Respirology (Carlton, Vic.) 17, 590–600

8. Schwartz, C., Bølling, A. K., and Carlsten, C. (2020) Controlled human exposures to wood smoke: a synthesis of the evidence. Particle and fibre toxicology 17, 49

9. Vogel, C. F. A., Kado, S. Y., Kobayashi, R., Liu, X., Wong, P., Na, K., Durbin, T., Okamoto, R. A., and Kado, N. Y. (2019) Inflammatory marker and aryl hydrocarbon receptor-dependent responses in human macrophages exposed to emissions from biodiesel fuels. Chemosphere 220, 993–1002

10. Traboulsi, H., Guerrina, N., Iu, M., Maysinger, D., Ariya, P., and Baglole, C. J. (2017) Inhaled Pollutants: The Molecular Scene behind Respiratory and Systemic Diseases Associated with Ultrafine Particulate Matter. International journal of molecular sciences 18

11. Øvrevik, J., Refsnes, M., Låg, M., Holme, J. A., and Schwarze, P. E. (2015) Activation of Proinflammatory Responses in Cells of the Airway Mucosa by Particulate Matter: Oxidant- and Non-Oxidant-Mediated Triggering Mechanisms. Biomolecules 5, 1399–1440

12. Weng, C. M., Wang, C. H., Lee, M. J., He, J. R., Huang, H. Y., Chao, M. W., Chung, K. F., and Kuo, H. P. (2018) Aryl hydrocarbon receptor activation by diesel exhaust particles mediates epithelium-derived cytokines expression in severe allergic asthma. Allergy 73, 2192–2204

13. Provoost, S., Maes, T., Pauwels, N. S., Vanden Berghe, T., Vandenabeele, P., Lambrecht, B. N., Joos, G. F., and Tournoy, K. G. (2011) NLRP3/caspase-1-independent IL-1beta production mediates diesel exhaust particle-induced pulmonary inflammation. J Immunol 187, 3331–3337

14. Tal, T. L., Simmons, S. O., Silbajoris, R., Dailey, L., Cho, S. H., Ramabhadran, R., Linak, W., Reed, W., Bromberg, P. A., and Samet, J. M. (2010) Differential transcriptional regulation of IL-8 expression by human airway epithelial cells exposed to diesel exhaust particles. Toxicology and applied pharmacology 243, 46–54

15. Afonina, I. S., Zhong, Z., Karin, M., and Beyaert, R. (2017) Limiting inflammation-the negative regulation of NF-κB and the NLRP3 inflammasome. Nat Immunol 18, 861–869

16. O’Driscoll, C. A., Gallo, M. E., Fechner, J. H., Schauer, J. J., and Mezrich, J. D. (2019) Real-world PM extracts differentially enhance Th17 differentiation and activate the aryl hydrocarbon receptor (AHR). Toxicology 414, 14–26

17. Larigot, L., Juricek, L., Dairou, J., and Coumoul, X. (2018) AhR signaling pathways and regulatory functions. Biochimie open 7, 1–9

18. Bock, K. W. (2018) From TCDD-mediated toxicity to searches of physiologic AHR functions. Biochem Pharmacol 155, 419–424

19. Wilson, C. L., and Safe, S. (1998) Mechanisms of ligand-induced aryl hydrocarbon receptor-mediated biochemical and toxic responses. Toxicologic pathology 26, 657–671

20. Rothhammer, V., and Quintana, F. J. (2019) The aryl hydrocarbon receptor: an environmental sensor integrating immune responses in health and disease. Nature Publishing Group

21. Rossner, P., Jr., Libalova, H., Vrbova, K., Cervena, T., Rossnerova, A., Elzeinova, F., Milcova, A., Novakova, Z., and Topinka, J. (2020) Genotoxicant exposure, activation of the aryl hydrocarbon receptor, and lipid peroxidation in cultured human alveolar type II A549 cells. Mutation research 853, 503173

22. Luo, Y. H., Kuo, Y. C., Tsai, M. H., Ho, C. C., Tsai, H. T., Hsu, C. Y., Chen, Y. C., and Lin, P. (2017) Interleukin-24 as a target cytokine of environmental aryl hydrocarbon receptor agonist exposure in the lung. Toxicology and applied pharmacology 324, 1–11

23. Bowman, D. M. J. S., Williamson, G. J., Abatzoglou, J. T., Kolden, C. A., Cochrane, M. A., and Smith, A. M. S. (2017) Human exposure and sensitivity to globally extreme wildfire events. Nature Ecology and Evolution 1, 58–58

24. Abatzoglou, J. T., and Williams, A. P. (2016) Impact of anthropogenic climate change on wildfire across western US forests. Proceedings of the National Academy of Sciences 113, 11770–11775

25. Ghio, A. J., Soukup, J. M., Dailey, L. A., Tong, H., Kesic, M. J., Budinger, G. R. S., and Mutlu, G. M. (2015) Wood Smoke Particle Sequesters Cell Iron to Impact a Biological Effect. Chemical research in toxicology 28, 2104–2111

26. Dilger, M., Orasche, J., Zimmermann, R., Paur, H. R., Diabaté, S., and Weiss, C. (2016) Toxicity of wood smoke particles in human A549 lung epithelial cells: the role of PAHs, soot and zinc. Archives of toxicology 90, 3029–3044

27. Arif, A. T., Maschowski, C., Garra, P., Garcia-Käufer, M., Petithory, T., Trouvé, G., Dieterlen, A., Mersch-Sundermann, V., Khanaqa, P., Nazarenko, I., Gminski, R., and Gieré, R. (2017) Cytotoxic and genotoxic responses of human lung cells to combustion smoke particles of Miscanthus straw, softwood and beech wood chips. Atmospheric environment (Oxford, England: 1994) 163, 138–154

28. Danielsen, P. H., Møller, P., Jensen, K. A., Sharma, A. K., Wallin, H., Bossi, R., Autrup, H., Mølhave, L., Ravanat, J. L., Briedé, J. J., De Kok, T. M., and Loft, S. (2011) Oxidative stress, DNA damage, and inflammation induced by ambient air and wood smoke particulate matter in human A549 and THP-1 cell lines. Chemical Research in Toxicology 24, 168–184

29. Kocbach, A., Herseth, J. I., Låg, M., Refsnes, M., and Schwarze, P. E. (2008) Particles from wood smoke and traffic induce differential pro-inflammatory response patterns in co-cultures. Toxicology and applied pharmacology 232, 317–326

30. Sasse, S. K., Gruca, M., Allen, M. A., Kadiyala, V., Song, T., Gally, F., Gupta, A., Pufall, M. A., Dowell, R. D., and Gerber, A. N. (2019) Nascent transcript analysis of glucocorticoid crosstalk with TNF defines primary and cooperative inflammatory repression. Genome Res 29, 1753–1765

31. Gally, F., Sasse, S. K., Kurche, J. S., Gruca, M. A., Cardwell, J. H., Okamoto, T., Chu, H. W., Hou, X., Poirion, O. B., Buchanan, J., Preissl, S., Ren, B., Colgan, S. P., Dowell, R. D., Yang, I. V., Schwartz, D. A., and Gerber, A. N. (2021) The MUC5B-associated variant rs35705950 resides within an enhancer subject to lineage- and disease-dependent epigenetic remodeling. JCI insight 6

32. Huang da, W., Sherman, B. T., and Lempicki, R. A. (2009) Systematic and integrative analysis of large gene lists using DAVID bioinformatics resources. Nat Protoc 4, 44–57

33. Corces, M. R., Trevino, A. E., Hamilton, E. G., Greenside, P. G., Sinnott-Armstrong, N. A., Vesuna, S., Satpathy, A. T., Rubin, A. J., Montine, K. S., Wu, B., Kathiria, A., Cho, S. W., Mumbach, M. R., Carter, A. C., Kasowski, M., Orloff, L. A., Risca, V. I., Kundaje, A., Khavari, P. A., Montine, T. J., Greenleaf, W. J., and Chang, H. Y. (2017) An improved ATAC-seq protocol reduces background and enables interrogation of frozen tissues. Nat Methods 14, 959–962

34. Rubin, J. D., Stanley, J. T., Sigauke, R. F., Levandowski, C. B., Maas, Z. L., Westfall, J., Taatjes, D. J., and Dowell, R. D. (2020) Transcription factor enrichment analysis (TFEA): Quantifying the activity of hundreds of transcription factors from a single experiment. bioRxiv, 2020.2001.2025.919738

35. Grant, C. E., Bailey, T. L., and Noble, W. S. (2011) FIMO: scanning for occurrences of a given motif. Bioinformatics 27, 1017–1018

36. Lambert, S. A., Jolma, A., Campitelli, L. F., Das, P. K., Yin, Y., Albu, M., Chen, X., Taipale, J., Hughes, T. R., and Weirauch, M. T. (2018) The Human Transcription Factors. Cell 175, 598–599

37. Sasse, S. K., Mailloux, C. M., Barczak, A. J., Wang, Q., Altonsy, M. O., Jain, M. K., Haldar, S. M., and Gerber, A. N. (2013) The glucocorticoid receptor and KLF15 regulate gene expression dynamics and integrate signals through feed-forward circuitry. Mol Cell Biol 33, 2104–2115

38. Altonsy, M. O., Sasse, S. K., Phang, T. L., and Gerber, A. N. (2014) Context-dependent cooperation between nuclear factor kappaB (NF-kappaB) and the glucocorticoid receptor at a TNFAIP3 intronic enhancer: a mechanism to maintain negative feedback control of inflammation. J Biol Chem 289, 8231–8239

39. Park, E. J., and Park, K. (2009) Induction of pro-inflammatory signals by 1-nitropyrene in cultured BEAS-2B cells. Toxicology letters 184, 126–133

40. Verstrepen, L., Carpentier, I., and Beyaert, R. (2014) The biology of A20-binding inhibitors of NF-kappaB activation (ABINs). Adv Exp Med Biol 809, 13–31

41. Grilli, A., Bengalli, R., Longhin, E., Capasso, L., Proverbio, M. C., Forcato, M., Bicciato, S., Gualtieri, M., Battaglia, C., and Camatini, M. (2018) Transcriptional profiling of human bronchial epithelial cell BEAS-2B exposed to diesel and biomass ultrafine particles. BMC Genomics 19, 302

42. Dukler, N., Booth, G. T., Huang, Y. F., Tippens, N., Waters, C. T., Danko, C. G., Lis, J. T., and Siepel, A. (2017) Nascent RNA sequencing reveals a dynamic global transcriptional response at genes and enhancers to the natural medicinal compound celastrol. Genome Res 27, 1816–1829

43. Azofeifa, J. G., Allen, M. A., Hendrix, J. R., Read, T., Rubin, J. D., and Dowell, R. D. (2018) Enhancer RNA profiling predicts transcription factor activity. Genome Res

44. Azofeifa, J., Allen, M., Lladser, M., and Dowell, R. (2016) An annotation agnostic algorithm for detecting nascent RNA transcripts in GRO-seq. IEEE/ACM Trans Comput Biol Bioinform

45. Thorvaldsdóttir, H., Robinson, J. T., and Mesirov, J. P. (2012) Integrative Genomics Viewer (IGV): high-performance genomics data visualization and exploration. Briefings in Bioinformatics 14, 178–192

46. Azofeifa, J. G., and Dowell, R. D. (2017) A generative model for the behavior of RNA polymerase. Bioinformatics 33, 227–234

47. Yates, P. R., Atherton, G. T., Deed, R. W., Norton, J. D., and Sharrocks, A. D. (1999) Id helix-loop-helix proteins inhibit nucleoprotein complex formation by the TCF ETS-domain transcription factors. Embo j 18, 968–976

48. Esnault, C., Gualdrini, F., Horswell, S., Kelly, G., Stewart, A., East, P., Matthews, N., and Treisman, R. (2017) ERK-Induced Activation of TCF Family of SRF Cofactors Initiates a Chromatin Modification Cascade Associated with Transcription. Mol Cell 65, 1081–1095.e1085

49. Buenrostro, J. D., Wu, B., Chang, H. Y., and Greenleaf, W. J. (2015) ATAC-seq: A Method for Assaying Chromatin Accessibility Genome-Wide. Current protocols in molecular biology 109, 21.29.21–29

50. Feng, J., Liu, T., Qin, B., Zhang, Y., and Liu, X. S. (2012) Identifying ChIP-seq enrichment using MACS. Nat Protoc 7, 1728–1740

51. Tripodi, I. J., Allen, M. A., and Dowell, R. D. (2018) Detecting Differential Transcription Factor Activity from ATAC-Seq Data. Molecules (Basel, Switzerland) 23

52. Cartharius, K., Frech, K., Grote, K., Klocke, B., Haltmeier, M., Klingenhoff, A., Frisch, M., Bayerlein, M., and Werner, T. (2005) MatInspector and beyond: promoter analysis based on transcription factor binding sites. Bioinformatics 21, 2933–2942

53. Zoch, A., Auchynnikava, T., Berrens, R. V., Kabayama, Y., Schöpp, T., Heep, M., Vasiliauskaitė, L., Pérez-Rico, Y. A., Cook, A. G., Shkumatava, A., Rappsilber, J., Allshire, R. C., and O’Carroll, D. (2020) SPOCD1 is an essential executor of piRNA-directed de novo DNA methylation. Nature 584, 635–639

54. Yang, S. Y., Ahmed, S., Satheesh, S. V., and Matthews, J. (2018) Genome-wide mapping and analysis of aryl hydrocarbon receptor (AHR)- and aryl hydrocarbon receptor repressor (AHRR)-binding sites in human breast cancer cells. Archives of toxicology 92, 225–240

55. Gomez, A., Bindesbøll, C., Satheesh, S. V., Grimaldi, G., Hutin, D., MacPherson, L., Ahmed, S., Tamblyn, L., Cho, T., Nebb, H. I., Moen, A., Anonsen, J. H., Grant, D. M., and Matthews, J. (2018) Characterization of TCDD-inducible poly-ADP-ribose polymerase (TIPARP/ARTD14) catalytic activity. Biochem J 475, 3827–3846

56. MacPherson, L., Ahmed, S., Tamblyn, L., Krutmann, J., Förster, I., Weighardt, H., and Matthews, J. (2014) Aryl hydrocarbon receptor repressor and TiPARP (ARTD14) use similar, but also distinct mechanisms to repress aryl hydrocarbon receptor signaling. International journal of molecular sciences 15, 7939–7957

57. Vu, Y. H., Hashimoto-Hachiya, A., Takemura, M., Yumine, A., Mitamura, Y., Nakahara, T., Furue, M., and Tsuji, G. (2020) IL-24 Negatively Regulates Keratinocyte Differentiation Induced by Tapinarof, an Aryl Hydrocarbon Receptor Modulator: Implication in the Treatment of Atopic Dermatitis. International journal of molecular sciences 21

58. Rao, L. Z., Wang, Y., Zhang, L., Wu, G., Zhang, L., Wang, F. X., Chen, L. M., Sun, F., Jia, S., Zhang, S., Yu, Q., Wei, J. H., Lei, H. R., Yuan, T., Li, J., Huang, X., Cheng, B., Zhao, J., Xu, Y., Mo, B. W., Wang, C. Y., and Zhang, H. (2020) IL-24 deficiency protects mice against bleomycin-induced pulmonary fibrosis by repressing IL-4-induced M2 program in macrophages. Cell Death Differ

59. Williams, A. E., Watt, J., Robertson, L. W., Gadupudi, G., Osborn, M. L., Soares, M. J., Iqbal, K., Pedersen, K. B., Shankar, K., Littleton, S., Maimone, C., Eti, N. A., Suva, L. J., and Ronis, M. J. J. (2020) Skeletal Toxicity of Coplanar Polychlorinated Biphenyl Congener 126 in the Rat Is Aryl Hydrocarbon Receptor Dependent. Toxicological sciences: an official journal of the Society of Toxicology 175, 113–125

60. Wójcik-Pszczoła, K., Jakieła, B., Plutecka, H., Koczurkiewicz, P., Madeja, Z., Michalik, M., and Sanak, M. (2018) Connective tissue growth factor regulates transition of primary bronchial fibroblasts to myofibroblasts in asthmatic subjects. Cytokine 102, 187–190

61. Eze, I. C., Jeong, A., Schaffner, E., Rezwan, F. I., Ghantous, A., Foraster, M., Vienneau, D., Kronenberg, F., Herceg, Z., Vineis, P., Brink, M., Wunderli, J. M., Schindler, C., Cajochen, C., Röösli, M., Holloway, J. W., Imboden, M., and Probst-Hensch, N. (2020) Genome-Wide DNA Methylation in Peripheral Blood and Long-Term Exposure to Source-Specific Transportation Noise and Air Pollution: The SAPALDIA Study. Environmental health perspectives 128, 67003

62. Rider, C. F., and Carlsten, C. (2019) Air pollution and DNA methylation: effects of exposure in humans. Clinical epigenetics 11, 131

63. Laufer, B. I., Gomez, J. A., Jianu, J. M., and LaSalle, J. M. (2021) Stable DNMT3L overexpression in SH-SY5Y neurons recreates a facet of the genome-wide Down syndrome DNA methylation signature. Epigenetics Chromatin 14, 13

64. Suhaimi, N. F., Jalaludin, J., and Abu Bakar, S. (2020) Deoxyribonucleic acid (DNA) methylation in children exposed to air pollution: a possible mechanism underlying respiratory health effects development. Reviews on environmental health

65. Martin, E. M., and Fry, R. C. (2018) Environmental Influences on the Epigenome: Exposure-Associated DNA Methylation in Human Populations. Annual review of public health 39, 309–333

66. Kim, D. I., Song, M. K., and Lee, K. (2021) Diesel Exhaust Particulates Enhances Susceptibility of LPS-Induced Acute Lung Injury through Upregulation of the IL-17 Cytokine-Derived TGF-β(1)/Collagen I Expression and Activation of NLRP3 Inflammasome Signaling in Mice. Biomolecules 11

