## Supporting Information Index for "Genomics-based deconvolution of multiplexed transcriptional responses to wood smoke particles defines rapid AHR signaling dynamics" (from another version of this preprint: version 1, 2021-02-24)

**Supplemental Materials**

Supplemental Table S1. qPCR primer sequences.

Supplemental File S1. PRO-seq pipeline and QC reports.

Supplemental Table S2. ChIP-qPCR primer sequences.

Supplemental Table S3. DESeq2 output for WSP-induced early-peak genes identified by PRO-seq.

Supplemental Table S4. DESeq2 output for WSP-induced early-plateau genes identified by PRO-seq.

Supplemental Table S5. DESeq2 output for WSP-induced late genes identified by PRO-seq.

Supplemental Table S6. DAVID functional annotation analysis of early-peak genes.

Supplemental Table S7. DAVID functional annotation analysis of early-plateau genes.

Supplemental Table S8. DAVID functional annotation analysis of late genes.

Supplemental Table S9. DESeq2 output for early-peak bidirectional transcripts.

Supplemental Table S10. DESeq2 output for early-plateau bidirectional transcripts.

Supplemental Table S11. DESeq2 output for late bidirectional transcripts.

Supplemental Table S12. Genomic coordinates for all PRO-seq-identified bidirectional transcripts.

Supplemental Table S13. TFEA output for PRO-seq-identified enhancers.

Thoughts:	– should we include ATAC-seq QC and possibly MACS2 output?

- should we also include TFEA output for ATAC?
